# Variation in mutational (co)variances

**DOI:** 10.1101/2022.05.28.493854

**Authors:** François Mallard, Luke Noble, Charles F. Baer, Henrique Teotónio

## Abstract

Because of pleiotropy, mutations affect the expression and inheritance of multiple traits and are expected to determine the structure of standing genetic variation and phenotypic evolution. It is thus important to find if the **M** matrix, describing mutational (co)variances between traits, varies between genotypes. We here estimate the **M** matrix for six locomotion behavior traits in two genotypes of the nematode *Caenorhabditis elegans*. We find significant mutational variance along at least one phenotypic dimension of the **M** matrix, but its size and orientation was similar between genotypes. We then tested if the **M** matrices were similar to one **G** matrix describing the standing genetic (co)variances of a domesticated population derived by the hybridization of several genotypes and adapted to a lab defined environment for 140 generations. **M** and **G** are different in part because the genetic covariances caused by mutational pleiotropy in the two genotypes are smaller than those caused by standing linkage disequilibrium in the lab population. If generalized to other genotypes, these observations indicate that selection is unlikely to shape the evolution of the **M** matrix for locomotion behavior and suggests that the genetic restructuring due to the hybridization of *C. elegans* genotypes allows for selection in the lab on new phenotypic dimensions of locomotion behavior, phenotypic dimensions which are inaccessible to natural populations.

## 2 Introduction

The **G** matrix, the additive genetic (co)variance matrix summarizing how multiple traits are genetically structured and inherited from parents to offspring, provides both prospective and retrospective information about phenotypic evolution. Looking forward, the evolution of mean trait values over one generation, 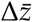, can be predicted from Lande’s equation (Lande, 1979): 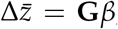, where *β* is the vector of directional selection gradients, cf. (Lande, 1976). Similarly, using the multivariate version of Bulmer’s equation, the evolution of trait variances depends on **G** and linkage disequilibrium (LD) generated by past selection (Bulmer, 1971; Tallis, 1987). Looking backward, the net selection gradients responsible for mean multivariate trait divergence between populations over multiple generations can be estimated from 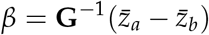 (Lande, 1979), where 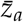 and 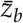are the vectors of trait means for the ancestral population *a* and the derived evolved population *b*. In this later case, the accuracy and precision of the inferences made depends on the stability of **G** over multiple generations (Arnold et al., 2001; Schluter, 1996; Shaw et al., 1995). However, and even in the absence of selection, **G** cannot be stable in the shortterm of tens to hundreds of generations, as mutation and genetic drift will change its orientation in unpredictable ways (Barton and Turelli, 2004; Mallard et al., 2019; Phillips and McGuigan, 2006; Phillips et al., 2001).

In the absence of selection, and assuming an infinitesimal model of trait inheritance, drift predictably removes genetic (co)variance from a diploid population at rate 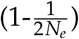 per generation (Barton et al., 2017; Lande, 1976; Lynch and Hill, 1986), where *N*_*e*_ is the effective population size. Mutation introduces genetic (co)variance into the population at rate **M** per generation, where the diagonal elements of **M** are the mutational variances, and the off-diagonals the mutational covariances between traits. In the long-term of mutation-drift equilibrium, *E*[**G**] = 2*N*_*e*_**M**, and the asymptotic rate of divergence in **G** between populations is 2**M** per generation (Felsenstein, 1988; Hansen and Martins, 1996; Lande, 1979; Lynch and Hill, 1986). Before reaching mutation-drift equilibrium, however, there is no stochastic theory to describe the expected distribution of **G**. One cannot predict the stability of **G** because all but the simplest deterministic models depend on the distribution of mutational effects, which for most traits is likely not normal, e.g. (Hayes and Goddard, 2001), and third and higher odd moments of the distribution might affect the evolution of **G** (Barton and Turelli, 1987, 1989; Johnson and Barton, 2005).

Examination of a simple deterministic model is nonetheless instructive as it suggests a way forward to better understand the evolution of **G**. Assuming that mutational effects are multivariate Gaussian and that selection is weak relative to recombination then (Lande, 1980):

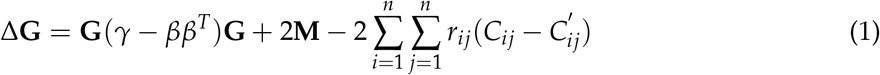

that is, changes in **G** depend on directional selection (*β*), stabilizing/disruptive selection on single traits and correlated selection between pairs of traits [diagonal and off-diagonal of *γ*, cf. Lande and Arnold (1983)], together with mutational input and output by recombination. The last term, represents the breakdown of genetic covariance resulting from LD between loci *i* and *j* with recombination rate *r*_*ij*_; with *C*_*ij*_ and 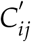 being the (co)variances resulting from associations between alleles on the same and different gametes, respectively. In the absence of other factors, selection will cause **G** to evolve to its expected value at a local point where fitness is maximized (Barton and Turelli, 1987; Cheverud, 1984; Jones et al., 2003; Lande, 1979). Mutation causes a buildup of LD, but there is no reason to expect that allelic effects of new mutations at different loci are correlated. Mutational covariance therefore reflects the underlying pleiotropic effects of new mutant alleles. Selection can maintain LD between combinations of alleles at different loci (Bulmer, 1976; Turelli, 1988), in which case genetic covariance results from both LD and pleiotropy. However, LD should be rapidly dissipated by recombination unless loci are very tightly linked or assortative mating is strong (Lande, 1980). As a consequence, Δ**G**=0 when the input of new genetic (co)variance by mutation offsets the (co)variance created by selection.

The deterministic model in equation 1 suggests that when a balance between mutation and selection is reached then the orientation of **M** should match that of **G**, and by extension that of the phenotypic divergence among taxa (**D**) (Lande, 1979). Two studies have found evidence for such congruence between **M** and **G** in *Drosophila* species wing shape (Dugand et al., 2021; Houle et al., 2017). Two other studies have found evidence for congruence between the orientations of **M** and **D**, one encompassing 40 million years of Drosophilid fly wing shape evolution (Houle et al., 2017), another encompassing 100 million years of Rhabditid nematode embryo size evolution (Farhadifar et al., 2015). Many other studies have further shown positive correlations between mutation and additive genetic variances (the diagonal elements of **M** and **G**) consistent with mutation-selection balance predictions, reviewed in Farhadifar et al. (2016). All these results are remarkable because **M** was measured in what effectively was a single genetic background of a single population from a single species. How can genetic and phenotypic evolution be predictable in the very long-term if **M** is bound to be subject to considerable sampling variance on the short-term because mutations are rare events? In addition, if mutation effects depend on genetic background (Pavličev and Cheverud, 2015), then correlated selection could lead to the evolution of **M** which in turn should affect standing levels in **G** and ultimately **D** (Hansen, 2006; Hermisson et al., 2003; Jones et al., 2007, 2014).

To understand the evolution of the **G** before reaching mutation-selection balance one should start by finding out about how variable is **M** between different genotypes, even if from a single population and species. Using six independent traits of locomotion behavior as a model in the hermaphroditic nematode *Caenorhabditis elegans*, we here characterize the **M** matrix in two genotypes and compare it with the **G** matrix of a lab adapted population with standing genetic variation. Comparison of the two **M** matrices between genotypes allow us to address the possibility for selection on standing levels of genetic covariances in locomotion behavior, while also indicating if genetic background effects are important for the expression of genetic variances of each component trait of locomotion behavior. Comparison of the **M** and **G** matrices allow us to question the relative role of pleiotropy and linkage disequilibrium in creating genetic covariances in locomotion behavior in a particular lab environment.

## 3 Methods

### 3.1 Archiving

Data for the domesticated lab population (A6140, see below) has been published in Mallard et al. (2019). All new data, modeling results (including **M** matrix estimates) and R code can be found in our github repository and will be archived in *Dryad*.*org* upon publication.

### 3.2 Experimental populations

To estimate **M** matrices on locomotion behavior we employed 250 generation mutation accumulation (MA) lines from two genotypes (N2 and PB306). The details of the derivation of these MA lines can be found in Baer et al. (2005) and Yeh et al. (2017). As at each generation only one hermaphrodite was passaged by selfing, we expect that most *de novo* mutations will be fixed within each lineage unless they are extremely deleterious (Teotónio et al., 2017).

We compared the **M** matrices with the **G** matrix of a lab population domesticated to standard lab conditions for 140 generations (called A6140). A6140 results from the hybridization of 16 founders (including N2 and PB306) for 33 generations followed by domestication to a standard lab environment at N=10^4^ census sizes and partial-outcrossing of 60%-80% for 140 4-day discrete and non-overlapping life-cycles (Mallard et al., 2019; Teotónio et al., 2012). From A6140, inbred lines were derived for 13-16 generations of single hermaphrodite selfing and the **G** matrix estimated as half the between inbred line differences (Mallard et al., 2019). In Mallard et al. (2019) we showed that this broad-sense **G** matrix is an adequate surrogate of the additive **G** matrix of the outbreeding A6140.

### 3.3 Locomotion behavior

Mutation accumulation lines were thawed from frozen stocks by blocks of 15 lines on 9cm Petri dishes and grown until exhaustion of food (*Escherichia coli* HT115). This occurred 2-3 generations after thawing, after which individuals were washed from plates in M9 buffer. Adults were removed by centrifugation, and two plates per line were seeded with 1000 larvae. Samples were then maintained for two complete generations in a common environment characterized by a 4-day non-overlapping life-cycle with extraction of embryos from adults done by “bleaching” and developmental synchronization of L1 larvae under starvation in M9 (Stiernagle, 1999). Growth from the L1 larval stage until adulthood was done in 9cm Petri dish plates with NGM-lite media and bacteria at a density of 1000. Full details about this standard lab environment, which was used during A6140 domestication as well, can be found in Teotónio et al. (2012).

At the assay generation (4-6 generations post-thaw), adults were phenotyped for locomotion behaviour at 72h post L1 stage seeding in single 9cm plates with NGM-lite media supplemented with bacteria, at a density of 1000. At the beginning of each assay we measured ambient temperature and humidity in the imaging room to control for their effects on locomotion. We phenotyped 54 and 62 mutation accumulation lines derived from the N2 and PB306 founders, respectively. 97 of these lines were included in two separate blocks, 1 line in 3 blocks and 18 lines were phenotyped only once. All thaw blocks contained the N2 MA ancestor, and 15 of 16 blocks contained the PB306 ancestor.

We imaged adults using the Multi-Worm Tracker, for a total of 246 plates [MWT version 1.3.0; Swierczek et al. (2011)]. We have followed the protocols detailed in Mallard et al. (2019) to image the worms and collect the data. Briefly, each movie contains about 1000 tracks of hermaphrodites (objects) with a mean duration of about 1 minute. Standardized to a common frame rate, we filtered and extracted the number and persistence of tracked objects per movie, and assigned movement states across consecutive frames as forward, still or backwards (assuming forward as the dominant direction of movement).

Modeling and estimation of the transitions rates per movie has also been detailed in (Mallard et al., 2019). We modelled the expected transition rates between forward, still and backward movement states with a continuous time Markov process. Given the constraint that self-transition rates equal the sum of the six transition rates between movement states, we consider that these six transition rates are independent. Log-likehood models were then specified in RStan (Stan Development Team (2018), R version 3.3.2, RStan version 2.15.1), which performs Bayesian inference using a Hamiltonian Monte Carlo sampling to calculate the posterior probability of the parameters given the observed data.

### 3.4 Mutational bias

We analysed the six independent mean transition rates in the MA lines using linear mixed models, for each founder MA genotype separately (N2 or PB306):

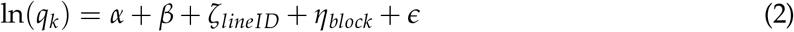

with *α* being the mean transition rate in the founder genotype before MA accumulation, *β* the mean difference between ancestral and MA lines, *ζ* ∼ 𝒩 (0, *σ*^2^) and *η* ∼ 𝒩 (0, *σ*^2^) the random effects of MA line identity and assay block; *ϵ* ∼ 𝒩 (0, *σ*^2^) is the residual error.

Differences of *β* to zero were tested with Likelihood Ratio Tests (LRT, which is approximately *χ*^2^ distributed), using the *anova* command in R and using as arguments two nested models containing or not the fixed effect. P values were corrected for multiple testing using the Benjamini-Hochberg method.

### 3.5 M-matrices

We estimated **M**-matrices separately for each founder genotype (N2 and PB306). The 6 transition rates *q*_*k*_ were fitted as a multivariate response variable *y* in the model:

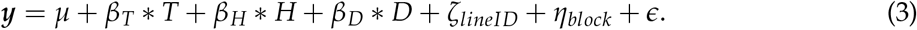

where *µ*_*k*_ is the general mean (intercept) of the *q*_*k*_ traits and *β*_*i*_ are fixed environmental effects of temperature (T), log density (D) and relative humidity (H) at the time of phenotyping for trait *q*_*k*_. *ζ, η* and *ϵ* as defined above for mutational bias. We then estimated a matrix of genetic (co)variance as half the line covariance matrix (*ζ*_*lineID*_).

Models were fit with the R package *MCMCglmm* (Hadfield, 2010). We defined priors as the phenotypic variances for each trait. Model convergence was verified by visual inspection of the posterior distributions and by ensuring that the autocorrelation remained below 0.05. We used 50,000 burn-in iterations, a thinning interval of 10 and a total of 500,000 MCMC iterations.

For each of the two **M** matrices, we constructed 1000 randomised **M** matrices to generate a null distribution. For this, we randomly shuffled MA line and block identities and fit the above equation in order to obtain 1,000 posterior distributions. As discussed in Walter et al. (2018), the significance of the posterior mean variance-covariance estimate is based on the overlap between the posterior null distribution of the posterior mean with the observed posterior mean. We consider that differences between the estimated empirical distributions can be inferred when their 80% credible intervals do not overlap Austin and Hux (2002).

### 3.6 Comparing M matrices

We performed eigen decomposition of each N2 or PB306 **M** matrices. The resulting first eigenvector usually contains most of the genetic variance, as measured by its eigenvalue *λ*, and can be thus be called *m*_*max*_ by analogy with the *g*_*max*_ of the **G** matrix of populations with standing genetic variation. Random sampling expectations for the six *λ* were obtained as above by shuffling block and MA line identity per N2 or PB306 genotype.

We compared the relative direction of the phenotypic dimension with more genetic variance between N2 and PB306 by computing the angle between their *m*_*max*_:

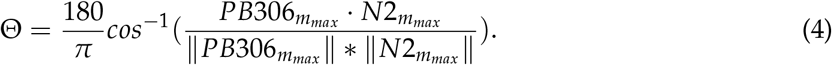

### 3.7 Comparing M and G matrices

The total amount of genetic (co)variances found in the N2 and PB306 **M** matrices and the A6140 **G** matrix depends on many factors, such as as the number of generations of mutation accumulation or the A6140 effective population size (see Introduction). **M** and **G** matrices therefore need to be standardized on a common scale. For this, we computed three new matrices with scaled total phenotypic variation by centering all transition rates to a mean of zero and dividing by the mean standard deviation of all the six transition rates. In this manner, the mean standard deviation of our six traits is one, though each of them has a standard deviation proportional to its initial value. We then ran the same models on each of the N2, PB306 and A6140 samples (equation 4). For the A6140 model we added a fixed effect of year of assay blocks.

To compare the size and shape of standardized **M** and **G** matrices we performed eigentensor analysis, as detailed in Aguirre et al. (2014); Hine et al. (2009). We used *MCMCglmm* for computation, while accounting for sampling variance as in Morrissey and Bonnet (2019). Briefly, eigentensors are 4-dimension objects describing overall genetic variation that can be decomposed into eigenvectors maximizing in orthogonal phenotypic space the amount of genetic differentiation between the three matrices. The first of these eigenvectors, usually explaining most genetic differentiation between populations, is called *e*_11_. Besides estimating the amount of genetic variance in this *e*_11_ vector (measured by its eigenvalue), we compared the angle between *g*_*max*_ of the A6140 with the *m*_*max*_ of N2 or PB306, as above in equation 4 by replacing *m*_*max*_ with *g*_*max*_. These angles reveal if the direction of the phenotypic dimension encompassing most genetic differentiation is aligned between A6140 with that of N2 or PB306.

## 4 Results

### 4.1 Mutation accumulation

To estimate the degree of mutation bias in locomotion behavior, we first compared the mean transition rates of the ancestral genotype founders to the mean of their respective mutation accumulation (MA) lines. We find that the two transition rates towards the still movement state (FS and BS) showed a significant decrease from the PB306 ancestor after 250 generations of mutation accumulation, while no difference was detected from N2 ancestor (Figure 1, Table 1).

**Table 1:** Mutational bias in transition rates. P values were obtained after a LRT following a *χ*^2^ distribution. Corrected P values for multiple comparisons were obtained with the Benjamini-Hochberg method

| Transition rate | Ancestor | P value | Corrected P value |
| --- | --- | --- | --- |
| SF | N2 | 0.31 | 0.52 |
| SB | N2 | 0.75 | 0.82 |
| FS | N2 | 0.14 | 0.40 |
| FB | N2 | 0.83 | 0.83 |
| BS | N2 | 0.16 | 0.40 |
| BF | N2 | 0.46 | 0.69 |
| SF | PB306 | 0.75 | 0.82 |
| SB | PB306 | 0.69 | 0.82 |
| FS | PB306 | 0.0036 | 0.022 |
| FB | PB306 | 0.073 | 0.29 |
| BS | PB306 | 0.00065 | 0.0078 |
| BF | PB306 | 0.29 | 0.53 |

**Figure 1:**
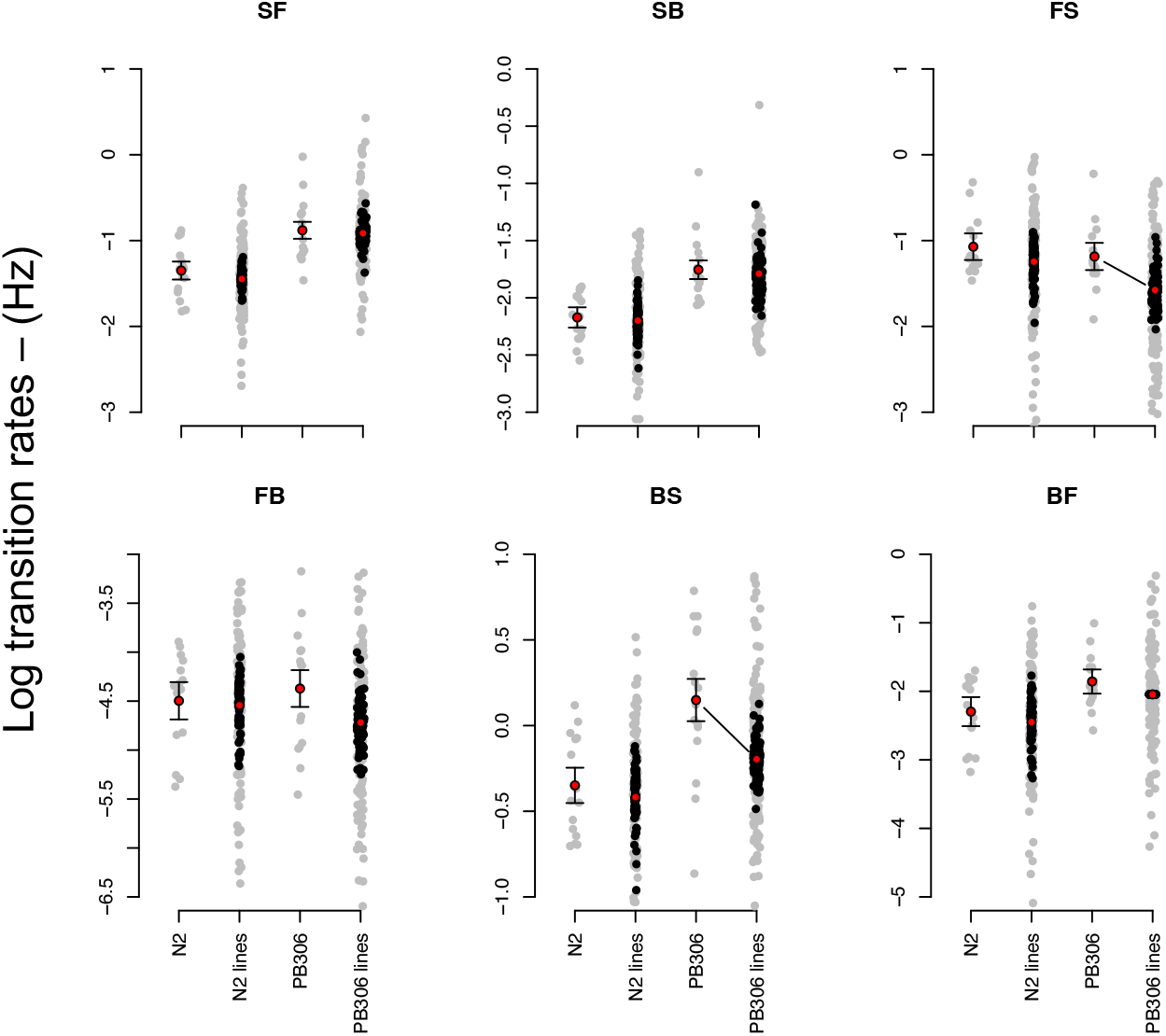
**A**. Mutational bias in locomotion behavior. Each plot show the transition rates between backward (B), forward (F) and still (S) movement states, with left to right letter ordering indi-cating the direction of movement. Red circles show the mean for the N2 and PB306 ancestor genotypes before and after mutation accumulation (MA). Grey dots show the uncorrected measurements and black the best linear unbiased predictors of the MA line means obtained from equation 2. Error bars in the ancestor genotypes are the standard error of the mean. In PB306, both FS and BS show a significant difference between ancestor and MA lines (lines, Table 1).

Next, we estimated the **M**-matrices of genetic variance-covariances created during the accumulation of new mutations during 250 generations (Figure 2). All genetic variances estimates are different from zero for both genotypes, but for the PB306 genotype only three of them are not expected from sampling alone. For genetic covariances between transition rates, most estimates of PB306 are also not different from what is expected from sampling, and in N2 covariances are significant but small.

**Figure 2:**
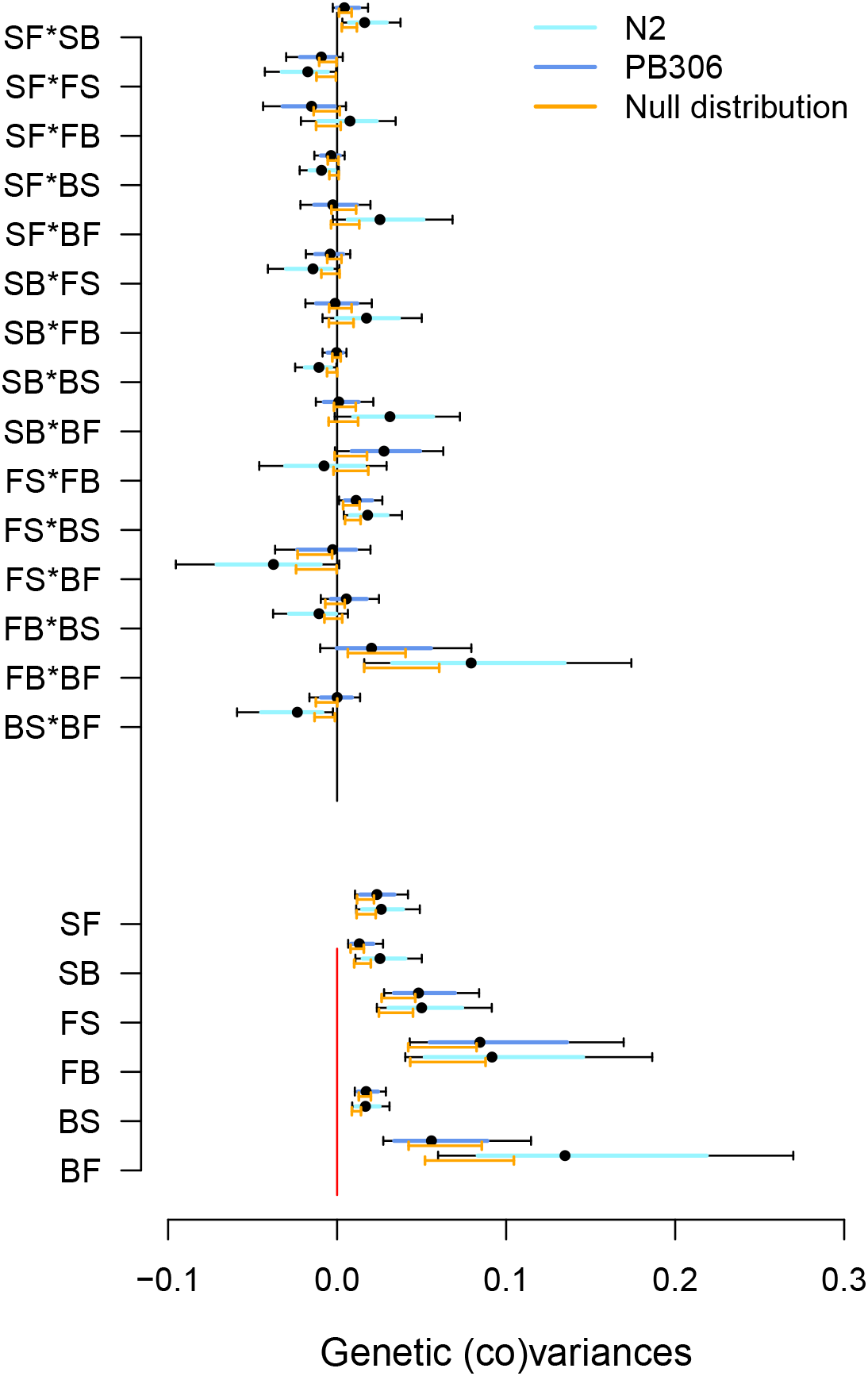
**A. M**-matrices for N2 and PB306 genotypes. The bottom six entries are the diagonal genetic variance estimates for each transition rate, while top 15 entries the off-diagonal genetic covariances estimates between transition rates. Null distributions were obtained by randomizing measurements and inbred line labels (see Methods). (Co)variances estimates are non-null if the posterior mean (black dot) is outside of the 95% CI of the distribution of randomized posterior means (orange).

We find, however, that the total amount of genetic variance in the **M** matrix does not differ between the N2 and PB306 genotypes (Figure 3A). Eigen decomposition further reveals that the phenotypic dimension encompassing most genetic variation, the first eigenvector *m*_*max*_, also does not differ between genotypes (Figure 3B). Because both **M** matrices are relatively unstructured by covariances calculating the angle between their respective *m*_*max*_ is uninformative (not shown).

**Figure 3:**
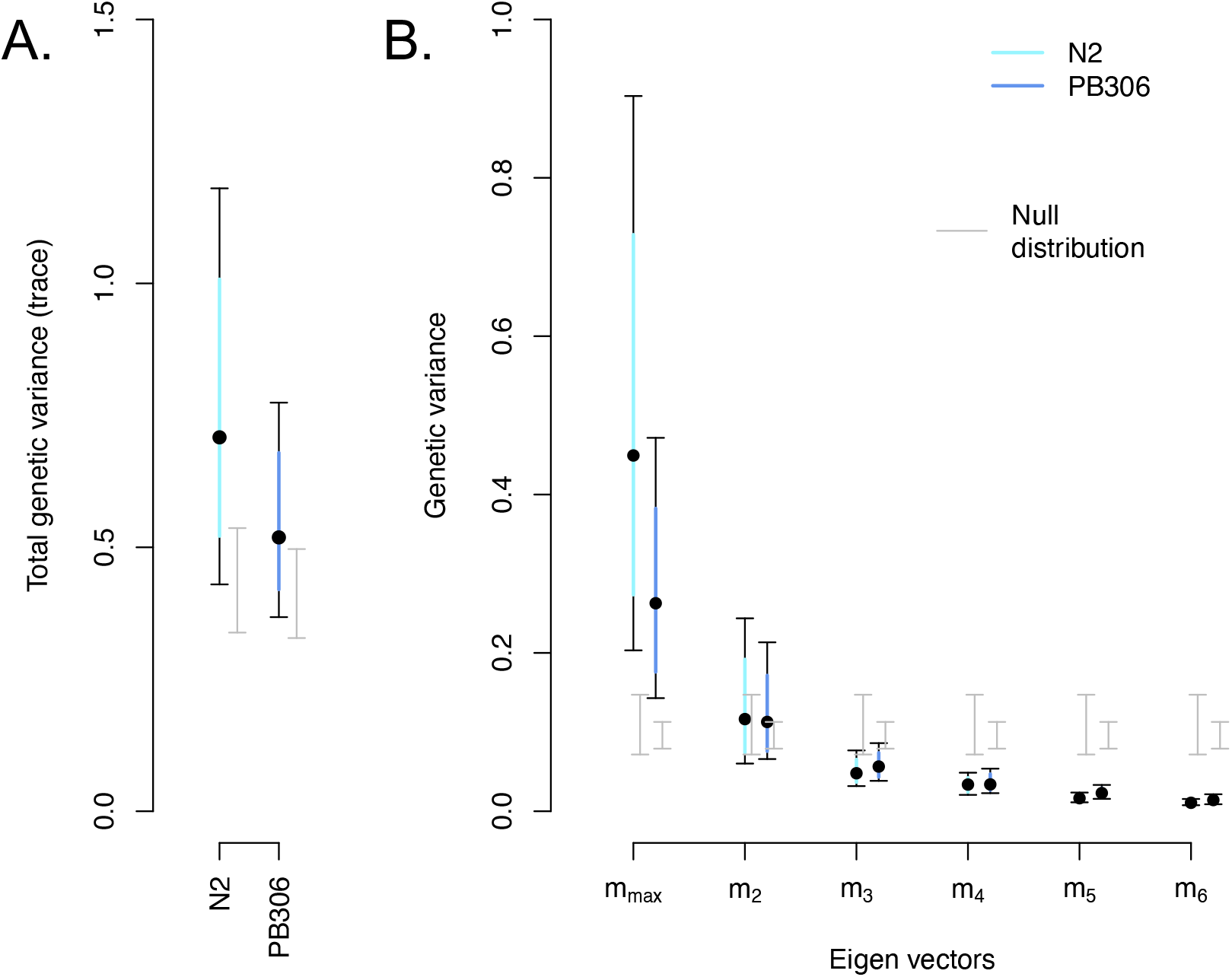
**M** matrix comparison between N2 and PB306. **A**. Shown is the total amount of genetic variance as measured by the trace of the **M** matrices. For both N2 and PB306, the total genetic variance is different from the randomized null distribution (grey, mean ± 95% CI), and there is no difference between N2 and PB306. **B**. Spectral decomposition of the **M**-matrices indicates that the dimension encompassing most genetic variance (measured by the *λ* eigenvalue), the first eigenvector *m*_*max*_, is different from the null distribution for N2 and PB306. N2 and PB306 do not differ in this *m*_*max*_ dimension.

### 4.2 Standing and mutation genetic variation

We compared the size and orientation of the N2 and PB306 **M** matrices with the **G** matrix of a lab domesticated population (A6140) containing standing genetic variation (see Methods). Detailed characterization of this **G** matrix has been presented in Mallard et al. (2019).

The **M** matrices of N2 and PB306 and the **G** matrix were first standardized to a common phenotypic scale (see Methods). We find that leaving the still movement state (SF and SB transition rates) contains more standing than mutational genetic variances (Figure 4), and that correspondingly the genetic covariances between these two transition rates with each other and with other transition rates differ between the lab domesticated population and N2 and PB306 genotypes.

**Figure 4:**
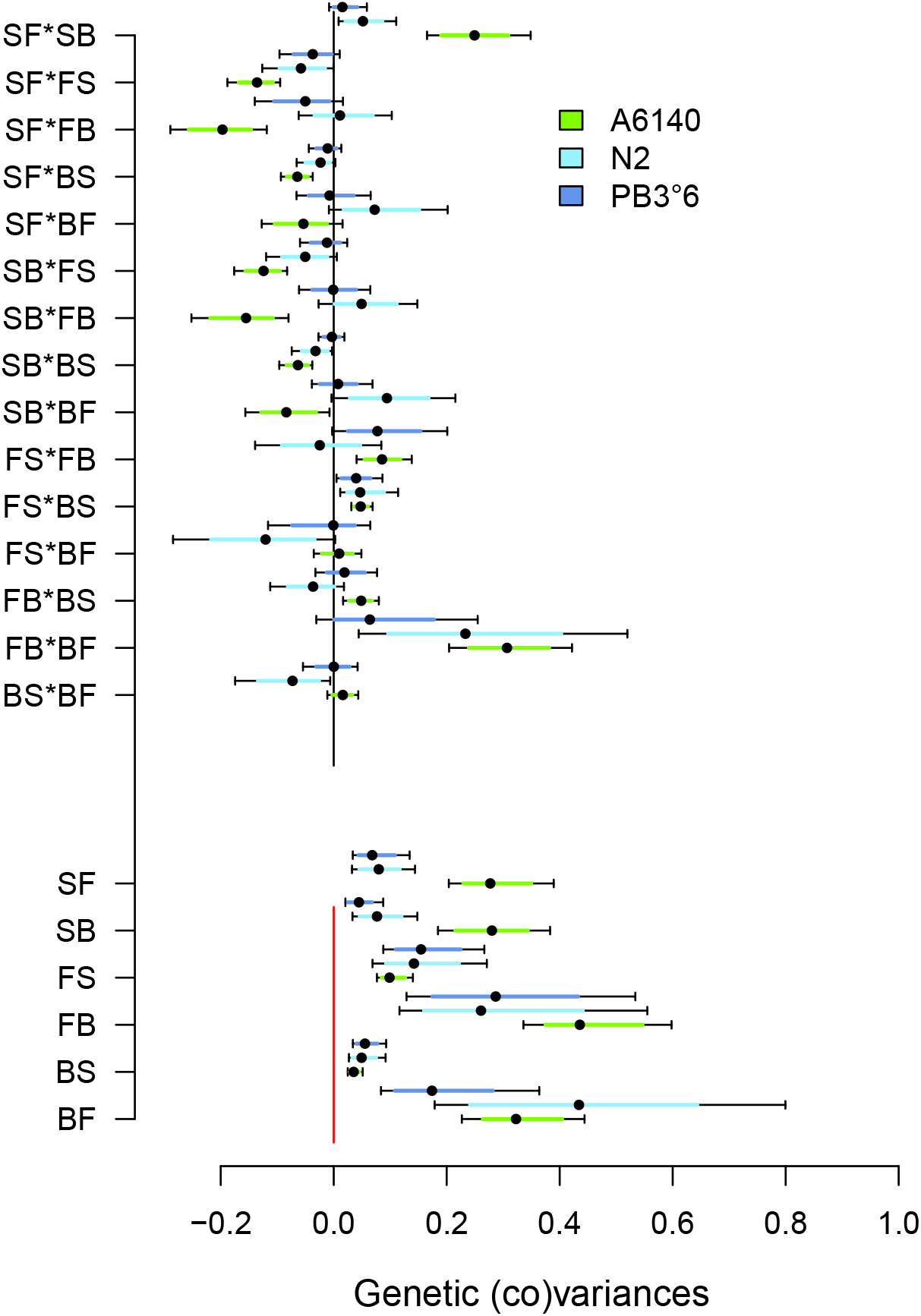
Standing and mutation genetic (co)variances for locomotion behavior. The bottom six entries are the diagonal genetic variance estimates for each transition rate, while top 15 entries the off-diagonal genetic covariances estimates between transition rates. Green for the lab adapted population (A6140), cyan for the N2 genotype, blue for the PB306 genotype. Dots show the mode of posterior distribution with bars being the 95% credible intervals. Distributions can be differentiated whenever their 80% credible intervals do not overlap (Austin and Hux, 2002). All estimates are standardized by on a common scale by dividing each genetic (co)variance by the total phenotypic variance in each population/genotype.

Spectral analysis of the three genetic (co)variance matrices reveals that only the first eigentensor is different from random expectations (Figure 5A), and that most genetic divergence is due to the lab adapted population (Figure 5B). Further decomposition of this eigentensor *E*_1_ shows that the lab adapted population is genetically differentiated from the N2 and PB306 genotypes only in the first eigenvector *e*_11_ (Figure 5C). The angle between the A6140 *g*_*max*_ and the two *m*_*max*_ vectors show a relatively high divergence between the main direction of genetic (co)variance between the G and M matrices even if the uncertainty remains high because of relatively low genetic structure in the M matrices.

**Figure 5:**
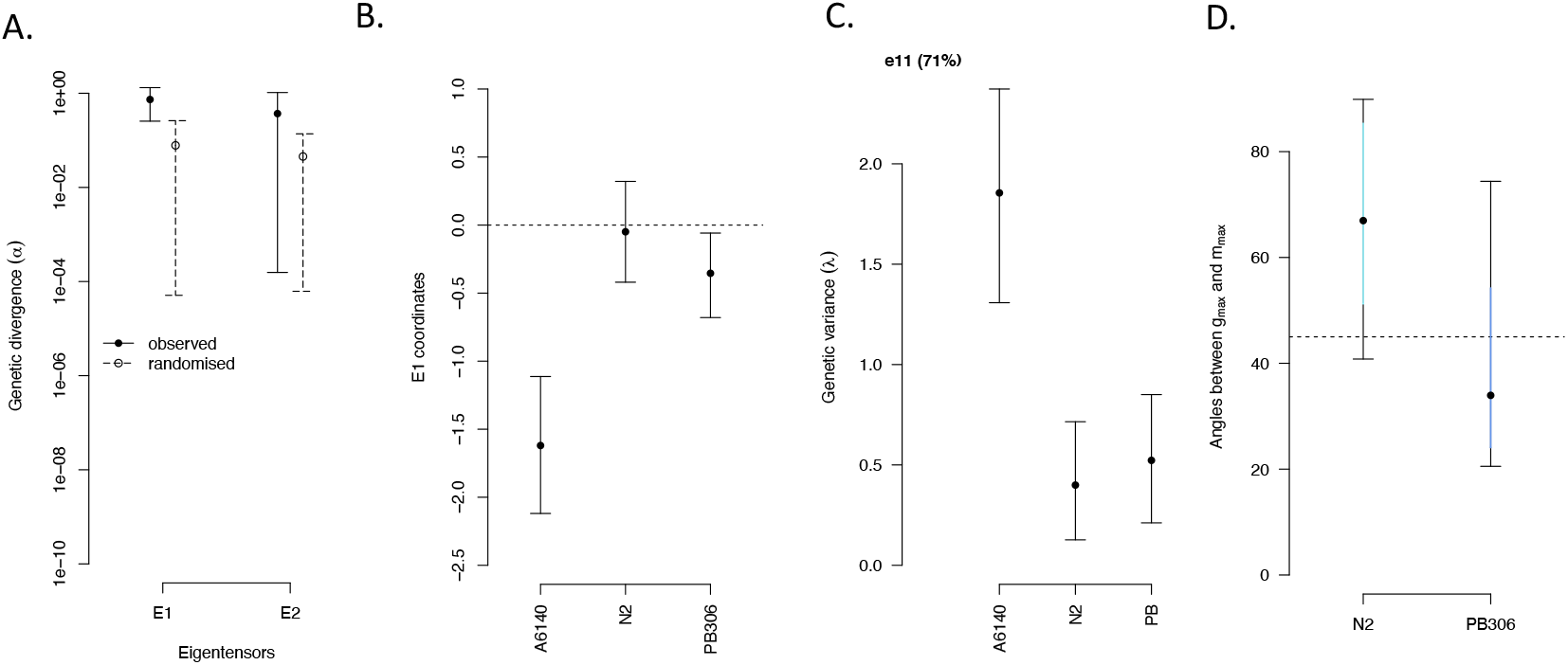
Standing versus mutation genetic (co)variances for locomotion behavior. **A**. Eigentensor decomposition of the three matrices (from Figure 4). The genetic variance explaining differences between matrices are shown as the mode and 95% credible interval of the posterior distributions of the first (*E*_1_) and second (*E*_2_) eigentensors, along with expected distributions by sampling (line and dashed, respectively). **B**. In the first eigentensor (*E*_1_), the coordinates of the three matrices. The lab adapted population (A6140) have the largest absolute values which shows that it drives most of the variation seen in Figure **??**A. **C**. The first eigenvector (*e*_11_) of the first eigentensor (*E*_1_) is the one where most genetic differences between the lab adapted population and the N2 and PB306 genotypes are found (71% of the variance found in E1). **D**. The angles between the *g*_*max*_ of the A6140 population with the *m*_*max*_ of the N2 or PB306 genotypes.

## 5 Discussion

We described the **M** matrix for locomotion behavior in two genotypes of *C. elegans*, and then compared them with the **G**-matrix of a lab adapted population. Our first finding is that the two **M** matrices of the N2 and PB306 genotypes are similar in size and orientation, though the N2 **M** matrix tends to be larger and more eccentric than the one from PB306. This result is in itself not too surprising given that both genotypes showed similar mutation rates and molecular spectra, except perhaps for the X chromosome, when a subset of MA lines from the two genotypes were whole-genome sequenced (Denver et al., 2012; Rajaei et al., 2021). Another study using the same two sets of MA lines used here showed that the mutational correlations between vulval and fitness-related traits, when significant, was similar in sign and magnitude among genotypes (Braendle et al., 2010). For both N2 and PB306, there is little evidence for pleiotropy because the genetic covariances between transition rates are much smaller than the genetic variances of each transition rate. These observations are consistent with those of studies of neural, muscular and metabolic function in (artificial) mutants in N2 showing that the direction of movement can be easily decoupled from activity (De Bono and Maricq, 2005; Zhen and Samuel, 2015). We also find, nonetheless, differential mutational bias between the two genotypes because for PB306 transitions rates between forward-still (FS), forward-backwards (FB) and backwards-still (BS) are consistently lower among the MA lines than the unmutated ancestor, while no such pattern is observed in the N2 genotype. Mutational bias could be interpreted to reveal epistasis (Halligan and Keightley, 2009; Jones et al., 2014; Saxena et al., 2018; Schweizer and Wagner, 2020), as the distribution of phenotypic effects depends on genetic background. However, with just two genotypes, and because MA lines carry a small and heterogeneous sample of mutations between them, the phenotypic effect distribution of mutations is barely sampled and any firm conclusion regarding epistasis remains unsupported (Jasmin and Lenormand, 2016; Keightley et al., 2000; Peters and Keightley, 2000).

By whole-genome sequencing many of the lines employed to estimate the **G** matrix of the lab population, as well as their 16 founders genotypes, we have previously reported that an exceptional high rate of *de novo* mutations appeared and were maintained at higher frequencies than expected with drift (Noble et al., 2021). Therefore, the result that is perhaps most surprising is that the **G** matrix of the lab adapted population was different from the **M** matrices. The two kinds of **M** and **G** matrices are different mostly due to more genetic covariation in the lab population between traits related to worm activity and leaving the still state. The simplest explanation for a difference between **M** and **G** is that other **M** matrices could be found in some of the other 14 genotypes that were founders of the lab populations, and some of them could be aligned with the **G** of the lab population. If so, the implication is that the **M** could evolve to orient itself with the selection surface(s) encountered in nature (Hansen, 2006; Hermisson et al., 2003; Jones et al., 2007, 2014). The only way forward to gain insights into this question is in obtaining more **M** matrices, from different genotypes, including those of the lab founders, and to compare them not only with the **G** of our lab population but also with the **G** matrix of the now more than 500 wild isolates in collection (Cook et al., 2017; Gilbert et al., 2022).

Assuming that all the **M** of the 16 founders of lab populations are similar, then differences between **M** and **G** could be explained by the selective purging of partially-dominant deleterious mutations during lab evolution. This is because to compare **M** and **G** we had to standardize them by their total respective phenotypic variation and thus an increase in genetic (co)variances for some traits is necessarily accompanied by a decrease in genetic (co)variance for some other traits, the expected pattern with purging. We do not see, however, a general trend for some traits showing less standing genetic (co)variance than mutational (co)variance; the exceptions being the standing genetic covariances between BS and BF, and that between FS and BF, which might be lower than the N2 mutational covariances. With a haploid base substitution mutation rate of 2.5×10^9^ (Saxena et al., 2018) and an effective population size of 10^3^ during lab evolution (Chelo and Teotónio, 2013), most of the mutations that we found segregating in the lab population should not have been observed (Noble et al., 2021). Purging of deleterious mutations is thus unlikely to have played a major role in shaping **G** during lab evolution.

Assuming an infinitesimal model of trait inheritance, elsewhere we showed that more loss of genetic variance than expected under drift occurred between the hybridization of the 16 isolates and initial lab evolution in phenotypic dimensions that were under stabilizing selection, a result indicating strong directional selection outside a small phenotypic region of locomotion behavior after adaptation (Mallard et al., 2019). In other words, there was effective stabilizing selection over a larger phenotypic space and inside the small phenotypic region there was stasis and the **G** matrix evolved as expected under neutrality. By definition, the **M** matrices reveal the genetic covariances due to pleiotropy while the **G** matrix should reveal genetic covariances due to both pleiotropy and linkage disequilibrium among the alleles at the relevant quantitative trait loci (QTL). The question thus becomes: how genetic covariances are maintained during our lab evolution? A difference between genetic covariances caused by pleiotropy or linkage disequilibrium under effective stabilizing selection in the lab could explain the difference between **M** and the lab domesticated **G** matrix.

*C. elegans* populations in nature are commonly found highly inbred and isogenic, due to a long history of predominant selfing, extensive selective sweeps and background selection (An-dersen et al., 2012; Cutter, 2006; Rockman et al., 2010). Hybridization of natural isolates leads to outbreeding depression (Chelo et al., 2013; Dolgin et al., 2007), in part due to the disruption of gene complexes (Gaertner et al., 2012; Seidel et al., 2008). We have previously described that lab adaptation not only involved maintenance of excess heterozygosity, due to overdominant loci interacting in a non-additive fashion (Chelo and Teotónio, 2013), but also that linkage disequilibrium, though much reduced from that found among founders, was still important for several genomic regions potentially encompassing many QTL (Chelo and Teotónio, 2013; Mallard et al., 2019; Noble et al., 2017). Hybridization of the 16 founders probably created the opportunity for the maintenance of linkage disequilibrium by selection (Bulmer, 1971; Tallis, 1987), in turn explaining why higher levels of genetic covariances are measured in the lab population than expected with mutation alone.

How do our results and interpretations compare with other studies, in other organisms and with different kinds of traits? To our knowledge, there has been only one other study in which the **M** matrix has been compared between genotypes within a species. Using a mutation accumulation design, Houle and Fierst (2013) compared **M** matrices for a set of up to 18 traits related to wing morphology between two of *Drosophila melanogaster* genotypes, genotypes that were known to have different mutation rates and molecular spectra (Schrider et al., 2013). When considering that alleles fixed within each MA line have additive effects, Houle and Fierst (2013) found that the two **M** matrices differ in both size and orientation. Interestingly, when considering also the heterozygous effects in crosses between MA lines, the two **M** matrices continued to differ but where dissimilar from additive **M** matrices. Directional dominance and/or epistasis was not detected, however, indicating that MA experiments are adequate, as a first approximation, to estimate the **M** matrix in organisms where inbreeding depression effects are common. In a later study, Houle et al. (2017) compared an “averaged” **M** matrix between the two genotypes with the orientation of the **G** matrix from a *D. melanogaster* population and with the divergence **D** matrix from many Drosophilid species spanning 40 million years, having found that they were all congruent in orientation with each other, if not in size. The authors concluded that mutation predicts genetic and phenotypic evolution. However, because only a single **M** matrix was used for prediction, it is unclear whether the **M** matrix can evolve to match the orientation of the selection surface as expected from theory (Jones et al., 2007, 2014).

In a more relevant study to compare with ours, Dugand et al. (2021) have recently estimated a single **M** matrix for five wing morphological traits but in the context of an evolving *D. serrata* outbred population for 14 generations, and from which a **G** matrix could be simultaneously estimated using a defined pedigreed experimental design (McGuigan et al., 2015). With this design, mutations appear, segregate and are fixed among many genotypes during evolution, and dominance and epistatic effects are explicitly accounted for. **M** and **G** are estimated on a common phenotypic scale, as there are no differential environmental effects, allowing then for the inference of selection in the long term of mutation-selection balance (Sztepanacz and Blows, 2017). Dugand et al. (2021) found that **M** and **G** were congruent for most phenotypic dimensions where there was significant standing genetic variation, except in one phenotypic dimension where less standing genetic variation was found than that expected for neutral traits with significant mutational variance. Not surprisingly for such a short-term experiment, selection on standing genetic diversity seemed to be more important than mutation.

Few studies that have attempted to measure how variation in mutational (co)variances determines standing genetic (co)variances and eventually phenotypic divergence between populations and species. While the study of Dugand et al. (2021) is outstanding in the excellence of design, it is unlikely to be much replicated because experiments and computing are prohibitively expensive. Mutation accumulation experiments have their own interpretation problems but they will be in a less expensive position to estimate **M** from specific genotypes. Given that these rare studies cannot be really compared, due to differences in design and analysis, little can be said about how the **M** matrix evolves, if it does, or about predicting phenotypic divergence from mutation in the short- or long-term. For locomotion behavior in the predominantly selfing *C. elegans*, hybridization of extant genotypes restructures genetic covariances in such a way that selection on the short-term at new phenotypic dimensions is possible when mutation has little influence.

## 6 Acknowledgments

We thank H. Gendrot and V. Pereira for help with worm handling, and M.A. Félix, L. Noble, P.C. Phillips and J. Picão-Osório for discussion. This work was partially supported by the Agence Nationale pour la Recherche (ANR-14-ACHN-0032-01, ANR-17-CE02-0017-01) to HT, and the National Institutes of Health (R01GM107227) to CB, and a Marie Curie fellowship (H2020-MSCA-IF-2017-798083) to LN.

## 7 Author contributions

Conceptualization FM, CB, HT; hardware and software implementation FM, TG; data acquisition and analysis FM, LN; funding acquisition CB, HT; project administration and resources CB, HT; writing, original draft FM, HT; review and editing LN, CB; correspondence FM, HT.

